# Gamma Music: A New Acoustic Stimulus for Gamma-frequency Auditory Steady-State Response

**DOI:** 10.1101/2023.08.17.552385

**Authors:** Yusuke Yokota, Kenta Tanaka, Chang Ming, Yasushi Naruse, Yasuhiko Imamura, Shinya Fujii

**Author notes:** Correspondence should be addressed to Y.Y.

## Abstract

A frequency range exceeding approximately 30 Hz, denoted as the gamma frequency range, is associated with various cognitive functions, consciousness, sensory integration, short-term memory, working memory, encoding and maintenance of episodic memory, and retrieval processes. In this study, we proposed a new form of gamma stimulation, called gamma music, combining 40 Hz auditory stimuli and music. This gamma music consists of drums, bass, and keyboard sounds, each containing a 40 Hz frequency oscillation. Since 40 Hz stimuli are known to induce an auditory steady-state response (ASSR), we used the 40 Hz power and phase locking index (PLI) as indices of neural activity during sound stimulation. We also recorded subjective ratings of each sound through a questionnaire using a visual analog scale. The gamma music, gamma drums, gamma bass, and gamma keyboard sounds showed significantly higher values in 40 Hz power and PLI compared to the control music without a 40 Hz oscillation. Particularly, the gamma keyboard sound showed a potential to induce strong ASSR, showing high values in these indices. In the subjective ratings, the gamma music, especially the gamma keyboard sound, received more relaxed, comfortable, preferred, pleasant, and natural impressions compared to the control music with conventional gamma stimulation. These results indicate that our proposed gamma music has potential as a new method for inducing ASSR. Particularly, the gamma keyboard sound proved to be an effective acoustic source for inducing a strong ASSR while preserving the comfortable and pleasant sensation of listening to music.

## Introduction

Human electroencephalogram (EEG) signals are designated as discrete frequency ranges, each correlating with distinct neural activity. Notably, a frequency range exceeding approximately 30 Hz, denoted as the gamma frequency range, is associated with various cognitive functions, consciousness, sensory integration, short-term memory, working memory, encoding and maintenance of episodic memory, and retrieval processes (Tallon-Baudry et al., 1998; Gruber et al., 2004; Herrmann et al., 2004; Jensen and Lisman, 2005; Mormann et al., 2005; Osipova et al., 2006; Fries et al., 2007; Lisman, 2010; Shirvalkar et al., 2010; Kucewicz et al., 2017; Griffiths et al., 2019). For instance, the amplitude of gamma frequency activity increases during the encoding of successfully memorized words compared to words that cannot be memorized (Gruber et al., 2004). In alternative instances, gamma frequency activity is detected during the preservation of visual stimuli in a short-term memory task (Tallon-Baudry et al., 1998), and the magnitude of gamma frequency activity is augmented commensurately with the number of items maintained in working memory (Howard et al., 2003; Van Vugt et al., 2010; Roux et al., 2012). Modulation of gamma frequency activity also has been reported in various neuropsychiatric disorders, including Alzheimer’s disease and schizophrenia (Herrmann and Demiralp, 2005; Mathalon and Sohal, 2015). Patients with Alzheimer’s disease exhibit reduced functional synchronization in gamma frequency activity during the resting state (Stam et al., 2002; Koenig et al., 2005) and protracted latencies in gamma frequency activity compared with healthy subjects (Başar et al., 2016). Similarly, during a series of cognitive tasks, including attention, memory, and object representation, gamma frequency activity in patients with schizophrenia is attenuated compared with healthy subjects (Shin et al., 2011). For instance, when presented with perceptually ambiguous images, also known as Mooney’s faces, gamma frequency activity in patients with schizophrenia is more attenuated than that in healthy subjects (Uhlhaas et al., 2006).

Numerous studies have reported that modulating neural activity within the brain during cognitive tasks can be observed as oscillations in the gamma frequency range. Conversely, in recent years, it has been reported that the presentation of stimuli possessing gamma frequency power can modulate cognitive processes within the brain. In a study involving mice, visual gamma frequency stimulation altered synaptic signaling and enhanced spatial learning and memory capabilities (Adaikkan et al., 2019). Similarly, visual and auditory gamma frequency stimulation has demonstrated the potential to improve Alzheimer’s disease biomarkers in mice, suggesting a possible alleviation of disease severity (Iaccarino et al., 2016; Martorell et al., 2019).

Presentation of periodic sensory stimuli amplifies the oscillatory activity corresponding to the stimulus frequency; this component is generally referred to as a steadystate evoked potential. The steady-state evoked potentials induced by auditory stimuli are commonly known as auditory steady-state responses (ASSR). ASSR is reportedly strongly evoked by a 40 Hz auditory stimulus (Galambos et al., 1981; Ross et al., 2000). The 40 Hz ASSR is modulated by various cognitive functions, such as human arousal levels (Griskova et al., 2007; Górska and Binder, 2019), attention levels (Skosnik et al., 2007; Gander et al., 2010; Voicikas et al., 2016), and workload (Yokota and Naruse, 2015; Yokota et al., 2017), as well as pathological symptoms such as schizophrenia (Kwon et al., 1999; O’Donnell et al., 2013; Thuné et al., 2016). Considering its impact on various cognitive functions, the magnitude of the ASSR elicited by the presentation of 40 Hz auditory stimuli holds potential as a biomarker for evaluating the degree of human cognitive function. The potential of 40 Hz stimulation to alleviate Alzheimer’s symptoms has been demonstrated previously. A 40 Hz amalgamated visual and auditory stimulation demonstrated diminished brain atrophy, attenuated loss of functional connectivity, and improved performance on an associative memory task (Chan et al., 2022). Since humans generally perceive simple 40 Hz auditory stimuli as unpleasant sounds (Fastl, 1990), this presents a considerable impediment to the social application of technologies that employ 40 Hz auditory stimuli. One potential approach to mitigate the unpleasantness associated with 40 Hz stimulation and enable humans to endure prolonged exposure is the integration of 40 Hz stimulation with music.

Listening to music is an emotive experience for many individuals, and it inherently functions as an enticing sensory stimulus (Zatorre and Salimpoor, 2013; Koelsch, 2014). One study reported that music-based interventions in patients with dementia improved their quality of life and emotional well-being (van der Steen et al., 2018). The integration of gamma stimulation technologies with music has been proposed and referred to as Gamma-MBI (music-based interventions) by Tichko et al., 2022. Tichko et al. (2022) argued that incorporating 40 Hz gamma frequency sounds into existing music presents challenges from the perspective of musical harmony because adding 40 Hz auditory stimuli to existing music can create an uncomfortable and unpleasant listening experience (see also Fastl, 1990). For example, in a study by Martorell et al. (2019), a sequence of tones repeating at 40 Hz (i.e., 1 ms long, 10 kHz tones played every 25 ms inter-tone intervals) was used as gamma auditory stimulus to ameliorate Alzheimer’s-associated pathology and enhance cognitive function. Since conventional gamma auditory stimulation is typically perceived as rough and unpleasant by listeners, it has been challenging to incorporate this stimulation into music while preserving the comfortable and pleasant sensation of listening (Tichko et al., 2022).

We conjectured that synthesizing the timbre of musical instruments, such as drum-kit sounds from 40 Hz oscillations—and modifying the amplitude envelope of musical instrument sounds, such as bass and keyboard sounds, into the gamma frequency range could solve this issue. Music can be divided into three fundamental elements: rhythm, melody, and harmony. Each of these elements has been reported to be associated with human perception, prediction, and emotional functioning (Vuust et al., 2022). We hypothesized that 40 Hz gamma auditory stimulation could be effectively integrated with these elements by creating a timbre of drum-kit sounds from gamma frequency tones and modifying the amplitude envelope of bass and keyboard sounds into the gamma frequency range. Accordingly, the gamma music used in this study was therefore composed of three musical instruments: a drum-kit, a bass guitar, and a keyboard, each representing the rhythm, melody, and harmony of music, respectively. The drum-kit sounds consisted of a kick drum, a snare drum, and a hi-hat sound. We synthesized each drum-kit sound from the gamma frequency tones, and termed them “gamma drums.” These three drum sounds were originally designed for this specific study by synthesizing them from a 40 Hz oscillatory component, with controlled envelopes to reproduce traditional drum sounds, along with noise sounds to enhance the percussive quality of the sounds (see Materials and Methods for details). The bass guitar and keyboard sounds provide melody in the lower registers and harmony in the higher registers, respectively. We supplemented a 40 Hz frequency component into the bass and keyboard sounds by using a Low-Frequency Oscillator (LFO) and termed these sounds “gamma bass” and “gamma keyboard,” respectively. As a control condition for gamma music, we used musical stimuli termed “control music” composed of drums, bass, and keyboard sounds that did not include a strong power at or around the 40 Hz frequency range. Additionally, as a third condition, we used the conventional gamma stimulation used by Martorell et al. (2019) and added it to the control music, terming it the “control music with conventional gamma stimulation (Conv. Gamma).”

We hypothesized that ASSR is elicited during the presentation of gamma music, gamma drums, gamma bass, gamma keyboard, and Conv. Gamma but not during the presentation of control music. We also hypothesized that the subjective ratings of gamma music, gamma drums, gamma bass, gamma keyboard, and control music are more comfortable and pleasant than those of Conv. Gamma. We used the power response and phase synchronization across trials (measured through the phase-locking index, PLI) for 40 Hz as indicators of ASSR. Additionally, we collected subjective ratings of the auditory stimuli from the participants using a Visual Analog Scale (VAS). Using these measurements of 40 Hz ASSR power, PLI, and subjective rating scales, we assessed the effectiveness of gamma music.

## Materials and Methods

### Participants

Seventeen individuals (nine men and eight women; age range: 20–41 years) participated in this study. All the participants had normal hearing. Prior to the experiment, the participants received a thorough explanation of the procedural details and provided written informed consent. This study was approved by the Shiba Palace Clinic Ethics Review Committee (Approval number: 152353_rn-35200) and conducted in accordance with the ethical principles outlined in the Declaration of Helsinki.

### Stimuli

One of the authors (KT) composed an original piece 196 second duration for this study using Ableton Live software (Ableton AG, Berlin, Germany). The piece was composed of MIDI signals and consisted of three musical-instrument tracks: a drum-kit, a bass guitar, and a keyboard. We utilized the “505 Core Kit,” “Basic Fine Line Bass,” and “Drifting Ambient Pad” preset sounds in Ableton Live to play each of the drums, bass, and keyboard MIDI tracks, respectively. Each audio track was output from Ableton Live and saved as a WAV audio file at a sampling frequency of 44,100 Hz. We referred to these output audio files as the “control drums,” “control bass,” and “control keyboard,” respectively. The “control music” was created by mixing these control drums, bass, and keyboard sounds.

Next, the control bass and keyboard sounds were loaded using MATLAB 2023 a software (MathWorks, Inc., Natick, Massachusetts, United States). We modified the amplitude envelope of each sound to 40 Hz using a custom-written program in MATLAB. We referred to these modified sounds as the “gamma bass” and “gamma keyboard,” respectively. As for the “gamma drums,” we created the bass drum, snare drum, and high-hat sounds from scratch using MATLAB. The bass drum sound had a carrier frequency of 40 Hz. The amplitude underwent decay and was supplemented with high-frequency noise during the attack phase to create a kick sound timbre. The snare sound has a carrier frequency of 40 Hz, and the amplitude underwent decay with the addition of bandpass noise to create a snare sound timbre. The high-hat sound was created by modifying the conventional gamma stimuli used by Martorell et al. (2019). The amplitude envelope of a train of gamma tones was decayed to create a high-hat timbre. The bass drum, snare drum, and high-hat sounds were saved as WAV audio files at a sampling frequency of 44,100 Hz in MATLAB and then loaded into Ableton Live to play the drum MIDI track. The output from Ableton Live was subsequently saved as a WAV audio file at a sampling frequency of 44,100 Hz. This drum-kit sound was referred to as the “gamma drums.” The “gamma music” was created by mixing these gamma drums, bass, and keyboard sounds. We also created conventional gamma tones using MATLAB software, based on the report by Martorell et al. (2019). We created a 1 ms long, 10 kHz tone played every 25 ms inter-tone interval, with a total duration of 196 seconds, and a sampling frequency of 44,100 Hz. The Conv. Gamma was created by mixing gamma tones with control music.

A total of six auditory stimuli were created: 1) gamma music, 2) gamma drums, 3) gamma bass, 4) gamma keyboard, 5) control music, and 6) Conv. Gamma. Perceptual loudness of all six stimuli was adjusted using the *integratedLoudness* function in MATLAB, performing loudness normalization in accordance with the EBU R 128 Standard with a target loudness level of – 23 LUFS. All stimuli were configured to fade in and out over the first and final five seconds. The waveforms and frequency characteristics of all the stimuli are shown in Figure 1. The music data are presented in Supplementary Data 1. During the experiment, all the stimuli were played as monaural sound sources in the first channel of the audio file. Additionally, an impulse signal synchronized with the gamma oscillation of the sound source was included every second as trigger information in the second channel. While the control music did not include a strong power at or around the 40 Hz frequency range, it was provided with trigger information at the same temporal timing as with the gamma music. This method was used for the PLI analysis, as described later in the study. The monaural sound source in the first channel was transformed into stereo and then delivered through headphones to both ears.

**Figure 1:**
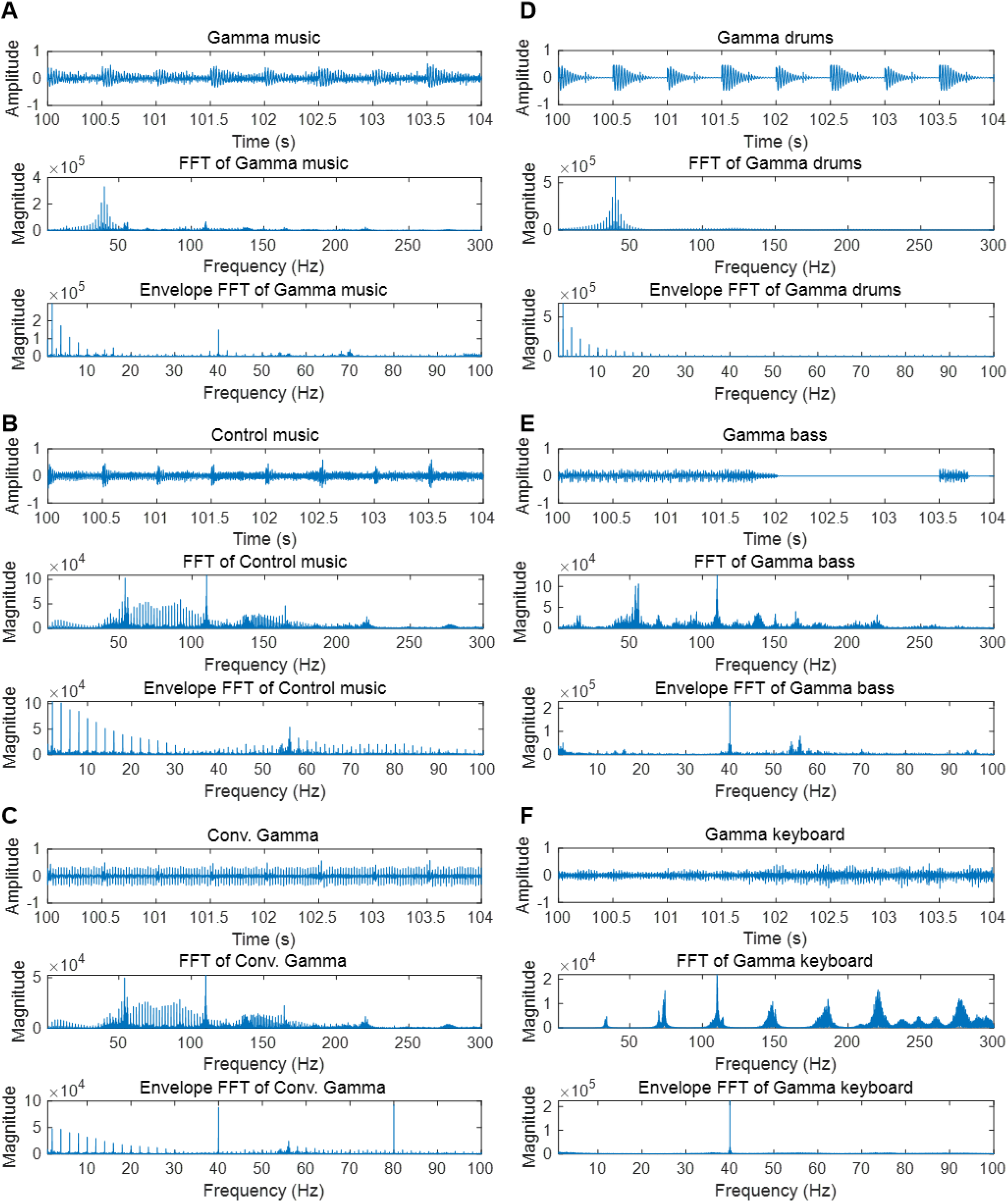
Partial waveform of each stimulus, the frequency response, and the frequency response of the auditory envelope. **A**. Gamma music **B**. Control music **C**. Control music with conventional gamma stimulation (Conv. Gamma) **D**. Gamma drums **E**. Gamma bass **F**. Gamma keyboard

### Experimental procedure

We presented six stimuli to the participants in random order. Participants were asked to relax their bodies and listen to music without any body movements. Participants were instructed to look at a fixation point displayed on the monitor in front of them to avoid unnecessary eye movements during the music presentation. After each music presentation, the participants self-assessed their mental relaxation and absorption levels (Rainville et al., 2002; Orozco Perez et al., 2020), as well as their valence and arousal levels (Russell, 1980). Furthermore, participants responded to several subjective rating items regarding the previously presented music. Table 1 presents all questionnaire items. All responses were measured using a VAS. After all recordings, the participants completed a self-report inventory using the Goldsmiths Musical Sophistication Index (GMSI) to assess individual differences in musical sophistication (Müllensiefen et al., 2014; Sadakata et al., 2022) and the Barcelona Music Reward Questionnaire (BMRQ) to assess the global sensitivity of the participants to music rewards (Mas-Herrero et al., 2013). All the subjective evaluations were conducted using the PsychoPy package (Peirce et al., 2019).

**Table 1:**
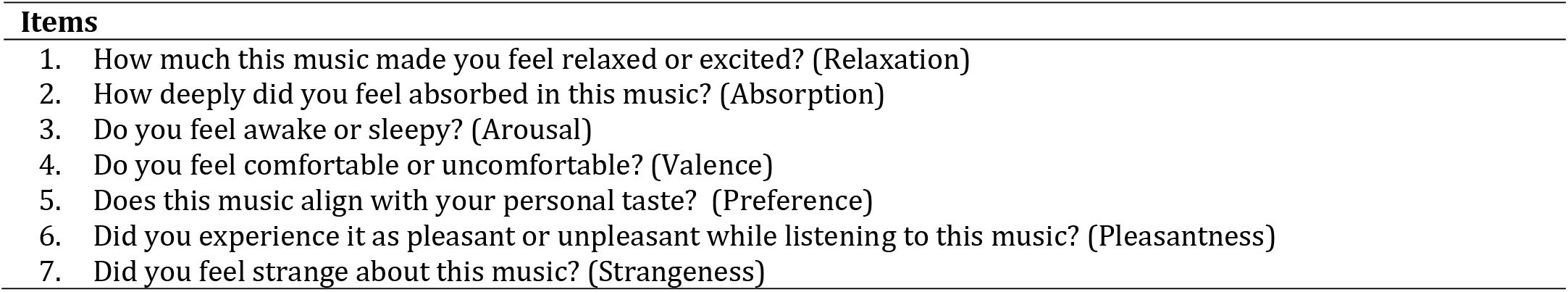
Subjective rating items.

### Experimental devices

EEG responses were recorded using a portable EEG device (Miyuki Giken, Bunkyō City, Tokyo, Japan; Polymate Mini AP108) with active electrodes placed at the forehead, FC3, FCz, and FC4, in accordance with the International 10–20 system. To obtain electrooculograms (EOGs), two electrodes were affixed to the superior and lateral aspects of each participant’s right eye. Moreover, to verify whether the components of the electrocardiogram (ECG) intermixed with those from the EEG electrodes, an electrode was affixed to the participant’s chest for ECG recording. All captured signals were referenced to the right mastoid, and the ground electrode was placed on the left. All electrode impedances were reduced to less than 50 kΩ. All the signals were sampled at 500 Hz.

All acoustic sources were produced via a USB audio interface (TASCAM, Santa Fe Springs, California, USA; US-4×4HR) and transmitted to the participants using headphones (Sony Corporation, Minato City, Tokyo, Japan; MDR-CD900ST). The delivery of all sounds was facilitated using a VLC media player (VideoLAN Organization, Paris, France).

### Data analysis

Data analyses were conducted using MATLAB 2023 a and R Studio version 2023.03.1. Build 446 (R Foundation for Statistical Computing, Vienna, Austria). Continuous EEG, EOGs, and ECG signals were digitally filtered using a finite-impulse response bandpass filter (1–50 Hz, order: 1500 Hz). Artifacts associated with eye movements were removed from the EEG data using independent component analysis.

We conducted a Fourier analysis of the continuous EEG data to convert it into the frequency domain for power analysis. Power was calculated as the square of the amplitude normalized using a factor of N, where N is the next power of two from the continuous EEG data. In this study, N was 131072 for all the experimental data. Subsequently, the power was transformed into decibel units using the calculation formula *20log10(power)*. Power was computed individually for each participant at each electrode and subsequently averaged across the forehead and fronto-central (FC3, FCz, and FC4) channel locations. Finally, the grand-average power was calculated by averaging the power values of the participants.

For PLI analysis, the continuous EEG data for each channel were segmented into 1000 ms epochs based on the auditory trigger that occurred every second in the continuous external input data. Epochs exceeding ± 40 μV on the forehead, FC3, FCz, and FC4 channels were removed from the analysis. A Fourier analysis was performed to convert the epochs into the frequency domain. PLI was calculated using the following formula:

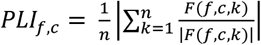

where *F* is the Fourier response, *f* is the frequency, *c* is the number of channels, *k* is the trial number and *n* is the number of trials. PLI was computed individually for each participant at each electrode and subsequently averaged across the forehead and fronto-central (FC3, FCz, and FC4) channel locations. Finally, the grand-average PLI was calculated by averaging the PLI values across participants.

### Statistical analysis

We used the Friedman test for subjective ratings. The factor under consideration was the stimulus, which was characterized by six auditory stimuli. Subsequent post-hoc tests were conducted using the Wilcoxon signed-rank test. The Holm method was used to correct the *p* values for multiple comparisons.

For the 40 Hz power and PLI data, we employed a linear mixed-effects model (LME) for statistical analysis. The stimuli (six auditory stimuli) and channel location (forehead or fronto-central) were defined as fixed effects. Participants were defined as having a random intercept. Our LMM model for Wilkinson notation was 40 Hz power or PLI values ∼ stimuli * channel location + (1 | Participant). The PLI values were subjected to a logit transformation before being applied to the LME. We performed LME analysis using the *lme4* and *lmerTest* packages in the R. A type III ANOVA test with the Kenward–Roger method was used to determine the significance of the fixed effects. For post-hoc tests, multiple comparisons were performed based on the fitted LME using the Kenward–Roger method. Tukey’s method was used to correct the *p* values for multiple comparisons. Post-hoc tests for the 40 Hz power and PLI were performed using *emmeans* package in the R.

Finally, we examined the potential impact of individual differences in musical sophistication and global sensitivity to music rewards on the magnitude of ASSR induced by sound stimuli. Specifically, we examined the correlation between individual GMSI and BMRQ scores and the power and PLI of the ASSR in the fronto-central region.

## Results

### Subjective ratings

The subjective ratings of the auditory stimuli are shown in Figure 2. The Friedman tests revealed significant differences in the stimuli for relaxation (*χ*^*2*^ = 24.8; *p* < 0.00015; *η*^*2*^= 0.29), valence (*χ*^*2*^ = 20.1; *p* = 0.0012; *η*^*2*^ = 0.24), preference (*χ*^*2*^ = 12.9; *p* < 0.024; *η*^*2*^ = 0.15), pleasantness (*χ*^*2*^ = 21.0; *p* = 0.0008; *η*^*2*^ = 0.26), and strangeness (*χ*^*2*^ = 23.4; *p* < 0.0003; *η*^*2*^= 0.29). However, no significant differences were found in the stimuli for absorption (*χ*^*2*^ = 2.45; *p* = 0.78; *η*^*2*^ = 0.029) and arousal (*χ*^*2*^ = 8.50; *p* < 0.13; *η*^*2*^ = 0.10).

**Figure 2:**
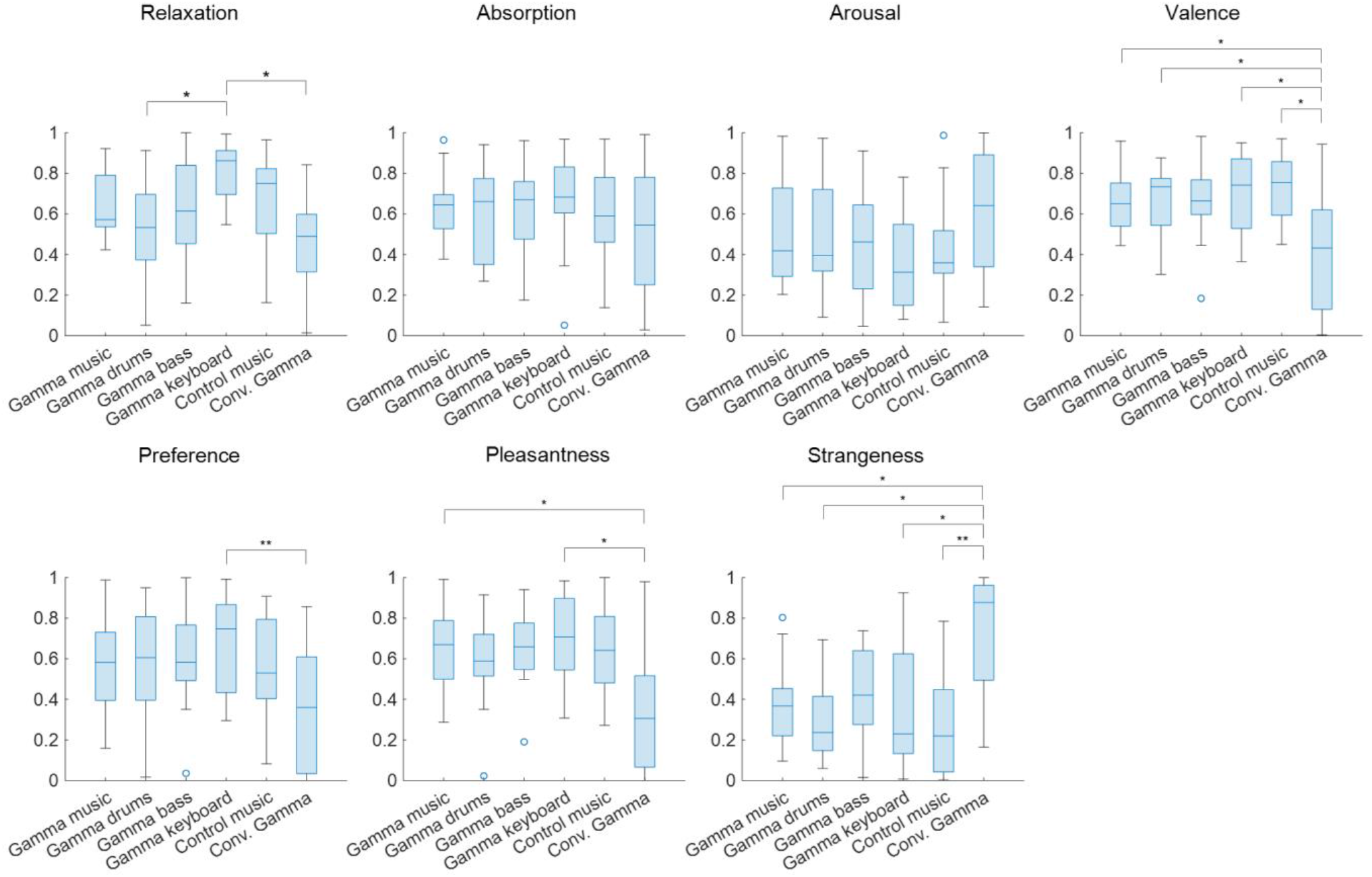
Box chart of subjective ratings for each of the sound stimuli. **: *p* < 0.01, *: *p* < 0.05

Subsequent multiple comparison tests of relaxation revealed a significant difference, such that the gamma keyboard sound was associated with a relaxed rating compared to the gamma drums sound (*Z* = 3.01; *p* = 0.036; *r* = 0.73) and the Conv. Gamma (*Z* = 3.95; *p* = 0.011; *r* = 0.96). Subsequent multiple comparison tests of valence revealed a significant difference, such that the Conv. Gamma was associated with uncomfortable ratings compared to the gamma music (*Z* = 3.27; *p* = 0.016 *r* = 0.79), the gamma drums sound (*Z* =3.01; *p* = 0.031; *r* = 0.73), the gamma keyboard sound (*Z* = 3.27; *p* = 0.016; *r* = 0.79), and the control music alone (*Z* = 3.14; *p* = 0.022; *r* = 0.76). Subsequent multiple comparison tests of preference revealed a significant difference, such that the Conv. Gamma was associated with an unpreferred rating compared to the gamma keyboard sound (*Z* = 3.48; *p* = 0.0076; *r* = 0.84). Subsequent multiple comparison tests of pleasantness revealed a significant difference, such that the Conv. Gamma was associated with unpleasant ratings compared to the gamma music (*Z* = 3.00; *p* = 0.038; *r* = 0.75) and the gamma keyboard sound (*Z* = 3.29; *p* = 0.015; *r* = 0.82). Subsequent multiple comparison tests of strangeness revealed a significant difference, such that the Conv. Gamma was associated with strangeness ratings compared to the gamma music (*Z* = 3.21; *p* = 0.019; *r* = 0.80), the gamma drums sound (*Z* =3.21; *p* = 0.018; *r* = 0.80), the gamma keyboard sound (*Z* = 3.14; *p* = 0.020; *r* = 0.79), and the control music alone (*Z* = 3.91; *p* = 0.0014; *r* = 0.98). These findings suggest that the participants found the Conv. Gamma to be the least unpreferred option. On the other hand, the gamma keyboard sound was the most favored, evoking sensations of a more relaxed, comfortable, preferred, pleasant, and natural impression.

The total GMSI score was 86.12 ± 17.48 (Mean ± SD), and the total BMRQ score was 84.12 ± 9.96 (Mean ± SD).

### EEG analysis

The grand-averaged power and PLI for each frequency in the forehead and fronto-central regions are shown in Figures 3 (A) and 4 (A). Clear peaks were observed in both the 40 Hz power and PLI within both the forehead and fronto-central regions. This indicates that the 40 Hz component included in the auditory stimuli induced ASSR. Figures 3 (B) and 4 (B) show box charts of the 40 Hz power and PLI for each stimulus in the forehead and fronto-central regions.

**Figure 3:**
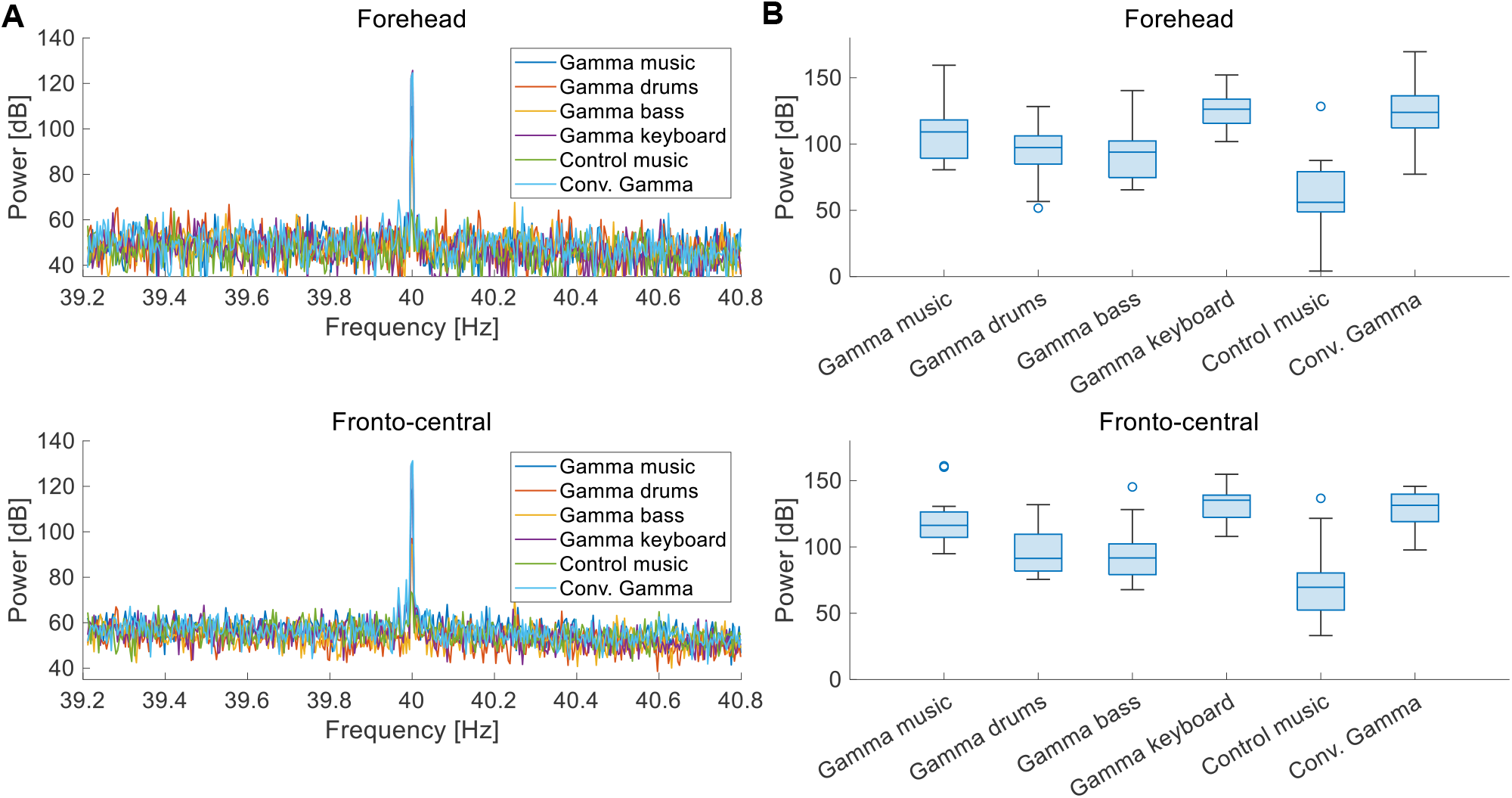
**A**. Grand-averaged power at the forehead and fronto-central regions of each sound stimulus. **B**. Box chart of 40 Hz power at the forehead and fronto-central regions of each sound stimuli.

**Figure 4:**
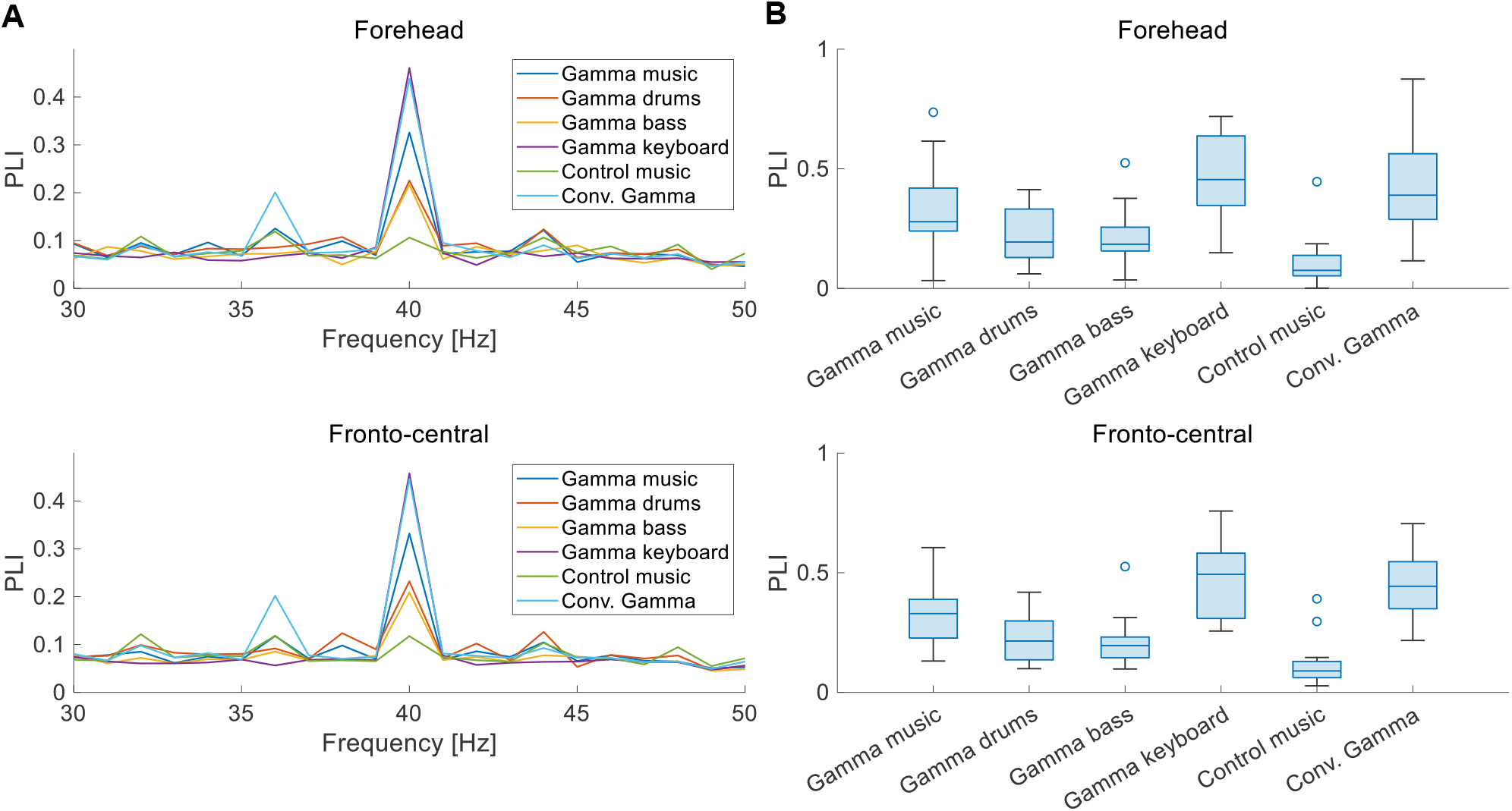
**A**. Grand-averaged PLI at the forehead and fronto-central regions of each sound stimulus. **B**. Box chart of 40 Hz PLI at the forehead and fronto-central regions of each sound stimuli.

The results of the statistical analyses for the 40 Hz power are summarized in Table 2. F-tests revealed the effects of both the stimuli (*F* (5, 176) = 35.3, *p* < 0.001) and channel location (*F* (5, 176) = 7.37, *p* < 0.0073). The results of the statistical analyses of the 40 Hz PLI are summarized in Table 3. F-tests revealed the effects of the stimuli (*F* (5, 176) = 30.0, *p* < 0.0001). Table 4 shows the results of the post-hoc analysis for a 40 Hz power. Table 5 shows the results of the post-hoc analysis for the 40 Hz PLI.

**Table 2:** Summary of statistical analyses for 40 Hz Power.

| Fixed effects | *F.value* | *d.f.* | *p val.* |
| --- | --- | --- | --- |
| Stimuli | 35.3 | (5, 176) | < 0.001** |
| Channel location | 7.37 | (1, 176) | 0.0073** |
| Stimuli : Channel location | 0.36 | (5, 176) | 0.87 |

**Table 3:**
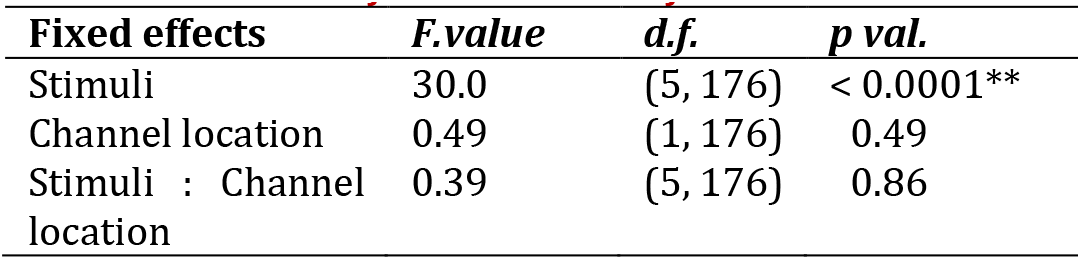
Summary of statistical analyses for 40 Hz PLI.

| Fixed effects | *F.value* | *d.f.* | *p val.* |
| --- | --- | --- | --- |
| Stimuli | 30.0 | (5, 176) | < 0.0001** |
| Channel location | 0.49 | (1, 176) | 0.49 |
| Stimuli : Channel location | 0.39 | (5, 176) | 0.86 |

**Table 4:**
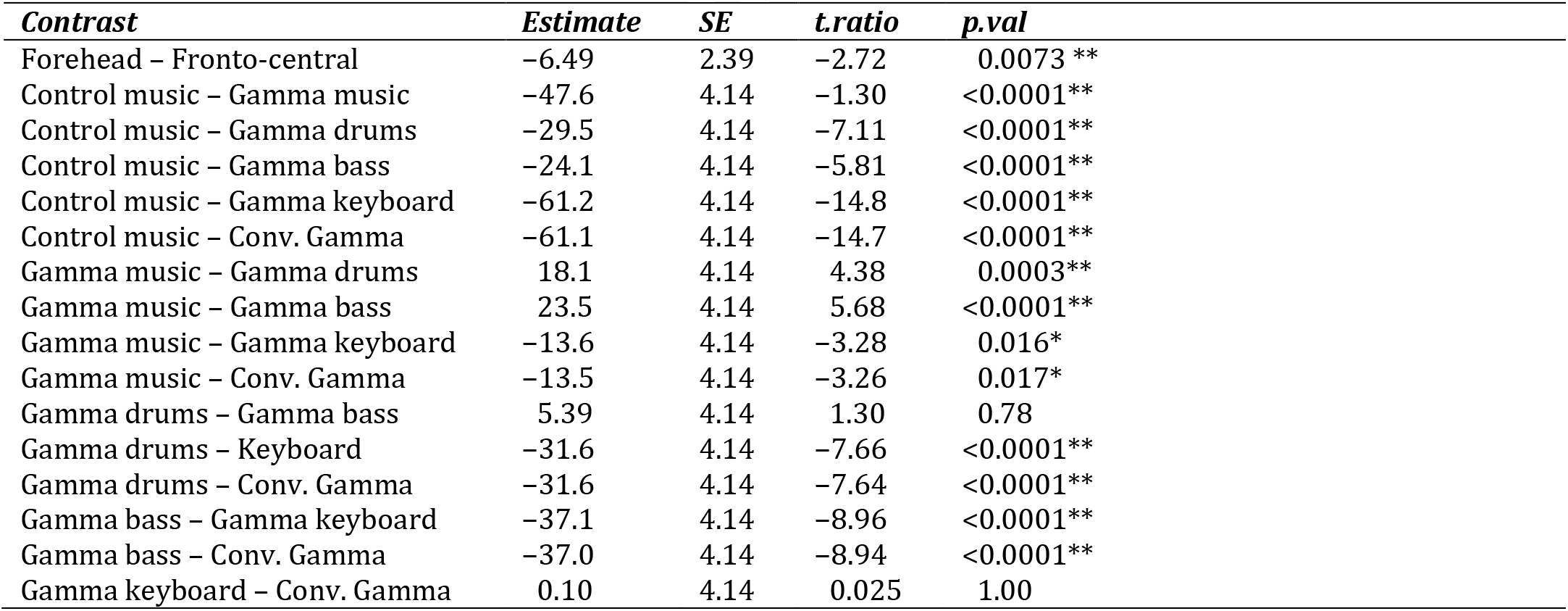
Summary of post-hoc test for 40 Hz Power.

| Contrast | Estimate | SE | *t.ratio* | *p.val* |
| --- | --- | --- | --- | --- |
| Forehead – Fronto-central | -6.49 | 2.39 | -2.72 | 0.0073 ** |
| Control music – Gamma music | -47.6 | 4.14 | -1.30 | <0.0001** |
| Control music – Gamma drums | -29.5 | 4.14 | -7.11 | <0.0001** |
| Control music – Gamma bass | -24.1 | 4.14 | -5.81 | <0.0001** |
| Control music – Gamma keyboard | -61.2 | 4.14 | -14.8 | <0.0001** |
| Control music – Conv. Gamma | -61.1 | 4.14 | -14.7 | <0.0001** |
| Gamma music – Gamma drums | 18.1 | 4.14 | 4.38 | 0.0003** |
| Gamma music – Gamma bass | 23.5 | 4.14 | 5.68 | <0.0001** |
| Gamma music – Gamma keyboard | -13.6 | 4.14 | -3.28 | 0.016* |
| Gamma music – Conv. Gamma | -13.5 | 4.14 | -3.26 | 0.017* |
| Gamma drums – Gamma bass | 5.39 | 4.14 | 1.30 | 0.78 |
| Gamma drums – Keyboard | -31.6 | 4.14 | -7.66 | <0.0001** |
| Gamma drums – Conv. Gamma | -31.6 | 4.14 | -7.64 | <0.0001** |
| Gamma bass – Gamma keyboard | -37.1 | 4.14 | -8.96 | <0.0001** |
| Gamma bass – Conv. Gamma | -37.0 | 4.14 | -8.94 | <0.0001** |
| Gamma keyboard – Conv. Gamma | 0.10 | 4.14 | 0.025 | 1.00 |

**Table 5:** Summary of post-hoc test for 40 Hz PLI.

| Contrast | Estimate | SE | *t.ratio* | *p.val* |
| --- | --- | --- | --- | --- |
| Control music – Gamma music | -1.65 | 0.17 | -9.99 | <0.0001** |
| Control music – Gamma drums | -1.11 | 0.17 | -6.75 | <0.0001** |
| Control music – Gamma bass | -1.03 | 0.17 | -6.23 | <0.0001** |
| Control music – Gamma keyboard | -2.27 | 0.17 | -13.7 | <0.0001** |
| Control music – Conv. Gamma | -2.20 | 0.17 | -13.4 | <0.0001** |
| Gamma music – Gamma drums | 0.53 | 0.17 | 3.24 | 0.0018* |
| Gamma music – Gamma bass | 0.62 | 0.17 | 3.76 | 0.0031** |
| Gamma music – Gamma keyboard | -0.62 | 0.17 | -3.75 | 0.0033** |
| Gamma music – Conv. Gamma | -0.56 | 0.17 | -3.40 | 0.011* |
| Gamma drums – Gamma bass | 0.086 | 0.17 | 0.52 | 1.00 |
| Gamma drums – Gamma keyboard | -1.15 | 0.17 | -6.99 | <0.0001** |
| Gamma drums – Conv. Gamma | -1.10 | 0.17 | -6.64 | <0.0001** |
| Gamma bass – Gamma keyboard | -1.24 | 0.17 | -7.51 | <0.0001** |
| Gamma bass – Conv. Gamma | -1.18 | 0.17 | -7.12 | <0.0001** |
| Gamma keyboard – Conv. Gamma | 0.058 | 0.17 | 0.35 | 1.00 |

The correlations between individual GMSI and BMRQ scores and the power and PLI of the ASSR in the frontocentral region for each sound stimulus are shown in Figure 5. No significant correlations were observed for any of the sound stimuli or their combinations (*p* > 0.05).

**Figure 5:**
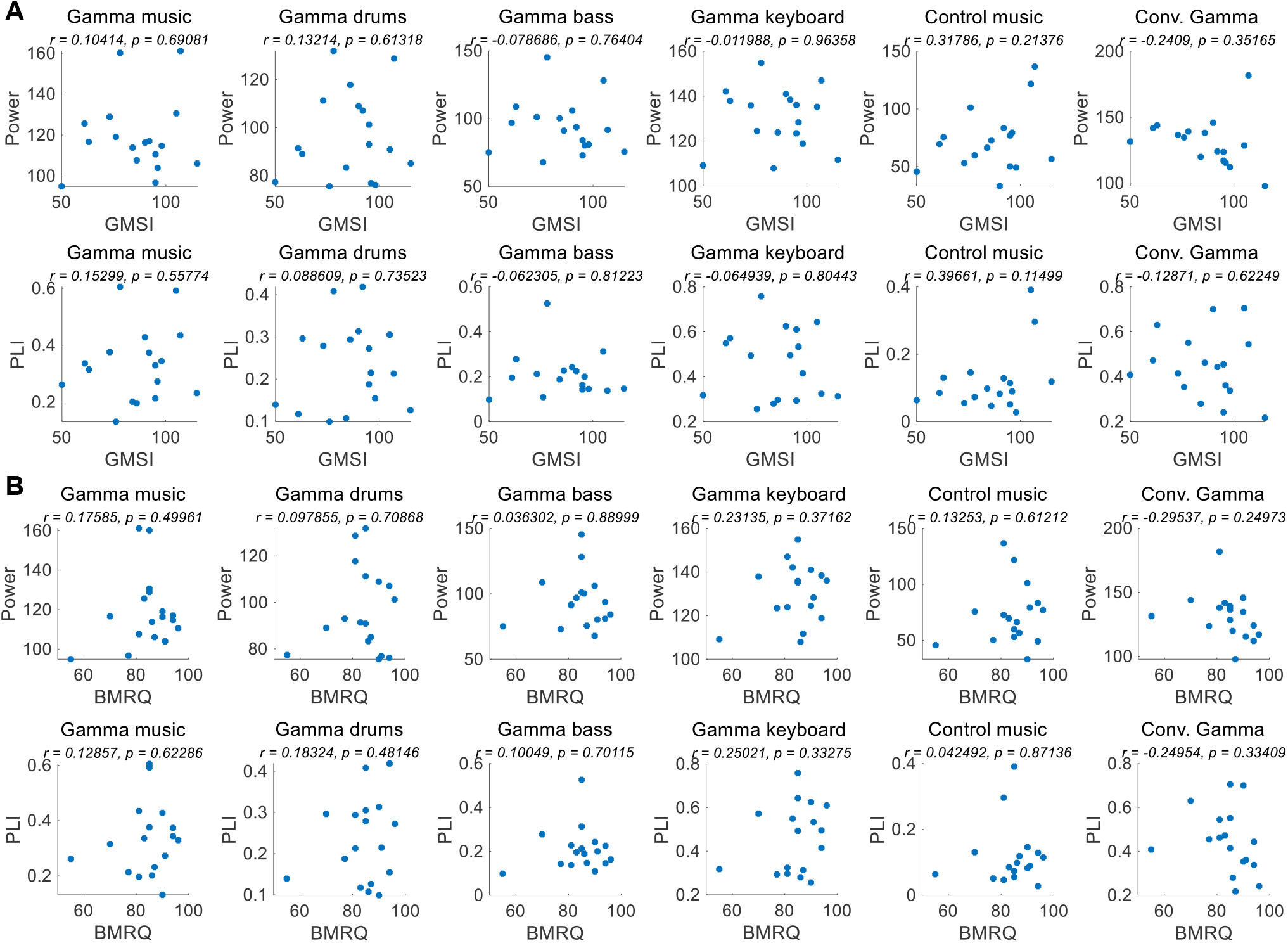
**A**. Correlation of ASSR indicators and GMSI score for each of the sound stimuli. **B**. Correlation of ASSR indicators and BMRQ score for each of the sound stimuli.

## Discussion

We propose “gamma music” as an innovative form of gamma stimulation that combines 40 Hz auditory stimuli and music. This gamma music was composed of drums, bass, and keyboard instruments, each of which included 40 Hz frequency oscillations. We used six stimuli: gamma music, the three individual instruments composing it, control music that did not include a strong power at or around the 40 Hz frequency range, and control music with conventional typical 40 Hz tones. As 40 Hz steady stimuli induce ASSR, we used the 40 Hz ASSR power and PLI as indicators of neural activity while listing the six stimuli. We also collected subjective ratings for each music stimulus through VAS questionnaires.

As a result of the experiment, the main effect of the fixed effects for stimuli was confirmed for both 40 Hz power and PLI. Gamma music, gamma drums, gamma bass, gamma keyboard, and Conv. Gamma showed significantly higher values at both 40 Hz power and PLI than the control music alone. This indicates that gamma music and the three gamma instruments strongly induced ASSR. In particular, the gamma keyboard sound demonstrated the highest value at both 40 Hz power and PLI, indicating its potential to strongly induce ASSR. The Conv. Gamma had a simple sequence of tones repeating at 40 Hz (i.e., 1 ms long, 10 kHz tones played every 25 ms inter-tone interval), commonly used in previous research (Martorell et al., 2019), and the degree of ASSR was comparable to that of gamma music and each gamma instrument created in this study. Notably, the Conv. Gamma elicited feelings of being unrelaxed, uncomfortable, non-preferred, unpleasant, and strange feelings compared to the other stimuli. This finding aligns with the assertion of Tichko et al. (2022) that solely adding a 40 Hz auditory stimulus to existing music presents difficulties from the perspective of musical pleasure. Conversely, the subjective ratings of the gamma music sound developed in this study were comparable to those of the control music. Among the stimuli, the gamma keyboard sound garnered the highest ratings for feelings of relaxation, comfort, preference, pleasantness, and naturalness. These results indicate that gamma music, particularly gamma keyboard sound can induce a strong ASSR while preserving the comfortable and pleasant sensation of listening to music.

Gamma music has various potential applications as a new methodology for inducing ASSR. A 40 Hz auditory stimulus is often perceived as an unpleasant sound by listeners (Fastl et al., 1990; Tichko et al., 2020). Therefore, prolonged stimulus presentation can potentially increase the burden on listeners. This is an issue in basic research investigating ASSR because the prolonged stimulus presentation time required to induce ASSR is generally an issue. In recent years, studies have suggested that gamma stimulation in mice and humans can modulate brain cognitive function, making the importance of gamma stimuli even more pronounced (Iaccarino et al., 2016; Martorell et al., 2019; Chan et al., 2022). Our gamma musical composition has the potential to resolve the discomfort experienced by participants during ASSR and clinical gamma interventions.

Our gamma music renders gamma stimulation a pleasant and comfortable experience for the participants. This holds significant value, especially if the ASSR can be reliably extracted, even when simple measurement methods are used. In this study, we measured EEG signals in both the fronto-central region, where the ASSR is typically strongly measured, and the forehead, where measurement is possible even with disposable electrodes with little interference from hair. In terms of 40 Hz power, significant main effects were observed for the fixed effects of the channel location, revealing higher power in the fronto-central region compared to the forehead. However, no significant main effect of the fixed effect on channel location was confirmed in the PLI. Therefore, while it is preferable to measure the fronto-central region when using 40 Hz power as an indicator of ASSR, it is adequate to solely measure the PLI values in the forehead when measuring the magnitude of ASSR induced by gamma music. In a previous study, PLI was confirmed to be a robust indicator, even against motion noise (Yokota et al., 2017), effectively reflecting the magnitude of the ASSR. Although EEG measurement in the fronto-central region is the most reliable method for evaluating the magnitude of ASSR, the measurement of PLI at the forehead using gamma music would be beneficial as a more straightforward and more comfortable method of gamma stimulation.

Why do gamma keyboard sounds elicit more significant power and PLI than gamma music? Gamma music, a composite of gamma drums, bass, and keyboard sounds, exhibited the second greatest power and PLI, with larger values than gamma drums and gamma bass sounds. There are two plausible explanations for these outcomes. The first is the attenuation of the loudness of gamma keyboard sounds. Gamma music contains a gamma keyboard sound. However, its loudness is lower than that of gamma keyboard sound alone. This discrepancy arises from the perceptual loudness that was adjusted throughout the experimental stimuli. Consequently, the loudness of each musical instrument sound included in the gamma music was quieter than that of each sound alone. While the gamma keyboard sound induced the largest ASSR power and PLI, the gamma drums and bass sounds induced a larger ASSR than the control music; however, the degree of these responses was relatively small. In gamma music, the loudness of the gamma drums and bass sounds becomes even smaller, resulting in a smaller ASSR than the gamma keyboard sound alone. The second possibility is that the time responses to the 40 Hz components of each instrument may vary, leading to the cancellation of the phases of each 40 Hz component. ASSR is a phenomenon in which brain activity settles into a steady-state owing to the convolution of a specific stimulus frequency. Therefore, when different stimuli are presented simultaneously, the neural responses to each stimulus may influence the others, resulting in a weaker ASSR. Therefore, even with stimuli at the same frequency, when multiple sensory stimulus presentations are used, it may be necessary to present stimuli and analyze the signals while considering their mutual influence.

In this study, we demonstrated that gamma music serves as an effective acoustic source for inducing a robust 40 Hz ASSR. In the future, rather than focusing solely on 40 Hz, we intend to identify the individual gamma frequency (IGF) that elicits the most substantial ASSR response for each individual, develop personalized gamma music based on these frequencies, and verify its efficacy. Cortical phase-locked activity in the gamma range varies significantly across individuals (Zaehle et al., 2010; Gransier et al., 2021). The ASSR to IGF is reported to correlate more robustly with individual cognitive indices than with the 40 Hz ASSR (Griškova-Bulanova et al., 2022). Consequently, the response to IGF may more accurately reflect the state of neural networks associated with information processing. Stimulating the IGF with gamma music may pave the way for more precise measurements of an individual’s ASSR and foster its application as an effective biomarker for cognitive evaluation.

## Conclusion

This study proposes a new form of gamma stimulation called gamma music, incorporating 40 Hz auditory stimuli into drums, bass, and keyboard instrument sounds. We showed that gamma music and each of the constituting gamma-sound instruments induced significantly higher values in 40 Hz ASSR power and PLI compared to control music that did not contain apparent 40 Hz frequency power. Notably, while the Conv. Gamma induced feelings of being unrelaxed, uncomfortable, non-preferred, unpleasant, and strange, gamma music, especially the gamma keyboard sound induced feelings of being relaxed, comfortable, preferred, pleasant, and natural in listeners. These results suggest that gamma music is a promising new method for inducing ASSR. In particular, the fact that our gamma sounds are pleasant to the human ear is beneficial for the long-term use of gamma stimulation. This music is expected to contribute to neuroscience research using ASSR and can be applied to gamma music-based interventions to enhance human cognitive functions.

## Acknowledgements

The authors would like to extend their gratitude to Keigo Yoshida and Taishin Emura for managing the recruitment of participants. We would like to thank Editage (www.editage.jp) for English language editing.

## Author contributions

Conceptualization: Y. N., Y. I., and S. F.

Data curation: Y. Y.

Formal analysis: Y. Y.

Funding acquisition: Y. I.

Investigation: Y. Y., C. M., and S. F.

Methodology: Y. Y., Y. N., Y. I., and S. F.

Project administration: Y. N., Y. I., and S. F.

Resources: Y. Y., K. T., and S. F.

Software: Y. Y., K. T., C. M., and S.F.

Supervision: Y. N., Y. I., and S. F.

Validation: Y. Y., C. M., and S. F.

Visualization: Y. Y. and S.F.

Writing – original draft: Y. Y., and S.F.

Writing – review & editing: Y. Y., K. T., C.M., Y. N., and S. F.

## Supplemental Data 1: Sound stimuli

